# SIMAIS: A swarm intelligence inspired biosensor for rapid protein detection

**DOI:** 10.1101/2025.10.31.685929

**Authors:** Jiahao Zheng, Bayinqiaoge, Hannah F. Schiff, Xin Wang, Taibo Wu, Xi Lu, Yuxin Zhang, Yi Wang, Shi-Yang Tang, Chengchen Zhang

## Abstract

Swarm intelligence, the collective behaviour of animals such as birds, fish, and ants, arises from simple local interactions and enables survival, efficient foraging, and decision-making without central control. Inspired by this principle, we aimed to harness swarm intelligence for biosensing applications. Conventional assays often struggle with limited sensitivity, long processing times, and the need for specialised instruments, restricting their use in decentralised healthcare. Here we developed a <u>s</u>warm intelligence of <u>m</u>icrobead-inspired, <u>a</u>rtificial intelligence (AI)-assisted, and <u>s</u>martphone-based (SIMAIS) biosensing platform that can transform invisible molecular recognition into visible, macroscale patterns. Millions of antibody-coated magnetic microbeads (MBs) organise into magnetically induced structures upon binding protein biomarkers, with the resulting patterns correlating directly to biomarker concentration. Using interferon-gamma (IFN-γ) as a model biomarker, we demonstrate that this approach delivers results within 10 minutes and achieves a tenfold increase in sensitivity compared to conventional lateral flow assays. Robust performance in human serum and urine confirms clinical applicability. By mimicking natural swarm behaviour, this platform introduces a new biosensing paradigm that bridges biology and engineering. Its versatility, speed, and ease of use highlight its potential for decentralised diagnostics, early disease detection, and broader applications in point-of-care healthcare.

## 1. Introduction

Swarm intelligence is a collective behaviour observed in groups of organisms arising from simple interactions among individuals and their environment^1-4^. In nature, many animals exhibit swarm intelligence, whereby collective behaviour enhances survivability, and facilitates complex decision-making in response to environmental changes^5-7^. This phenomenon arises from the self‐organised coordination of individuals following simple, decentralised rules, without reliance on a central leader^8,9^. For example, schools of fish synchronise to evade predators (**Figure 1a**). Swarm intelligence has inspired breakthroughs across diverse fields such as AI^10,11^, robotics^12-15^, optimisation algorithms^16-20^, and network systems^21-24^ to tackle computationally intensive and dynamic challenges. However, its transformative potential in healthcare remains largely underexplored.

**Figure 1.**
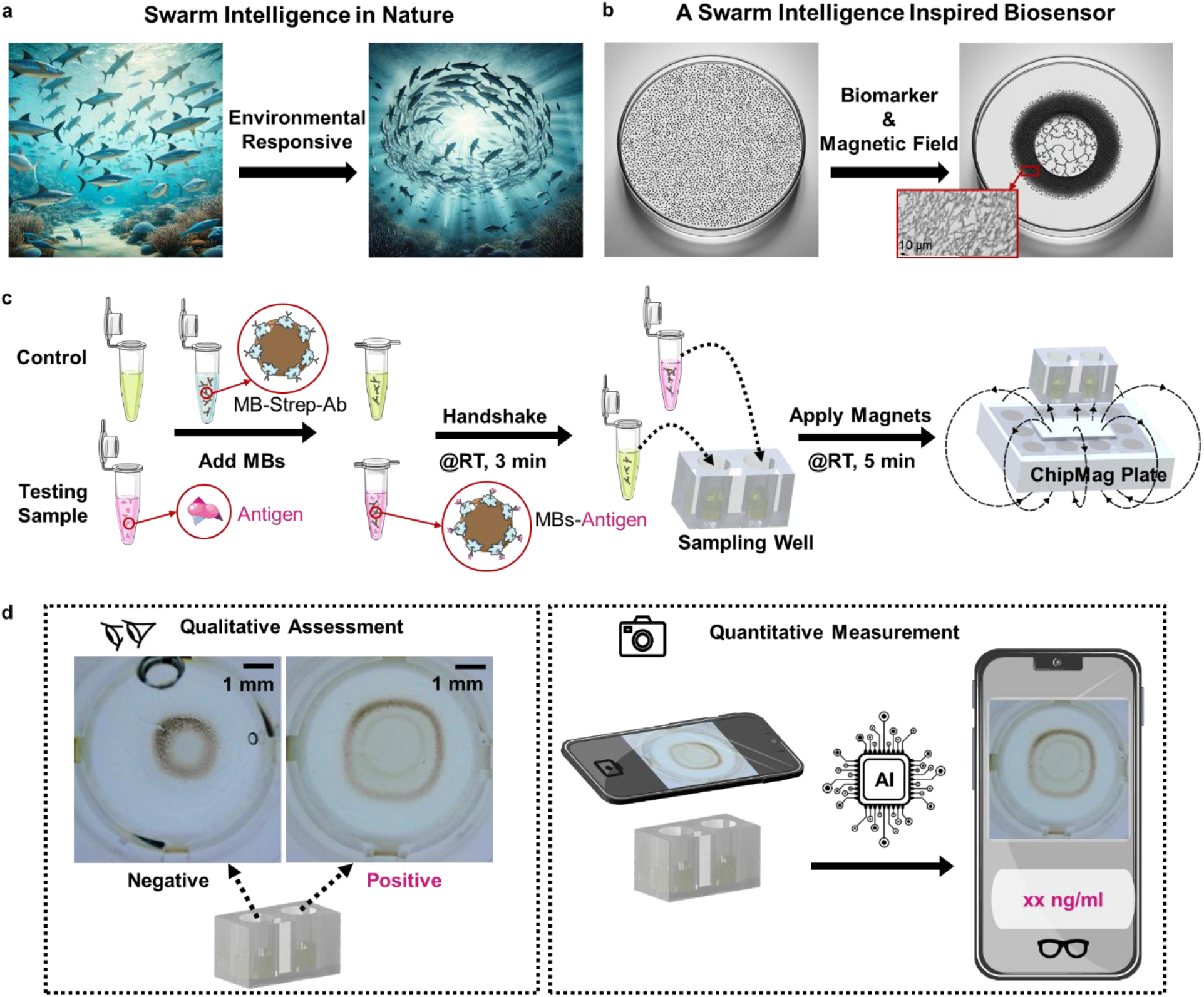
Overview of swarm intelligence inspired SIMAIS biosensor. a, Environmental responsive movements of fish schools indicating swarm intelligence in nature. b, Biomarker responsive deposition pattern of magnetic microbeads with external magnetic field applied. c, Schematic representation of the sample test workflow. d, Analysing approaches of SIMAIS biosensor test results (qualitatively assessable via naked eye and quantitatively measurable via AI model-deployed smartphone app).

Our study addresses this gap by introducing a biomarker-responsive biosensing platform that harnesses the principles of swarm intelligence. Specifically, we mimic the emergence of coordinated and distinct macroscale patterns in response to molecular cues – biomarkers – to achieve visually interpretable diagnostics, akin to natural swarm behaviour (**Figure 1b**). This platform employs antibody-coated magnetic microbeads (MBs) to selectively capture protein biomarkers. Upon exposure to varying analyte concentrations and an external magnetic field, the MBs dynamically organise into distinct, naked-eye-visible patterns that directly correlate with the presence and concentration of the target proteins. Enhanced with AI-enabled image analysis, the system enables rapid, sensitive and user-friendly on-site detection using a smartphone, significantly simplifying biomarker diagnostics by eliminating the need for bulky equipment and specialised training.

Beyond its conceptual novelty, the practical relevance of this platform lies in its ability to address a critical need in healthcare: efficient and accessible protein biomarker detection. Protein detection plays a crucial role in healthcare, as proteins serve as essential biomarkers for diagnosing, monitoring, and managing various diseases, from infections to chronic conditions and cancers^25-30^. Accurate and timely protein detection enables clinicians to identify disease states early, assess the progression of conditions, and make informed treatment decisions^31-34^. Currently, enzyme-linked immunosorbent assays (ELISA) are considered the gold standard due to their high specificity and sensitivity^35-37^. However, their long processing times and reliance on specialised equipment and trained personnel limit their suitability for rapid, decentralised settings such as pandemics and emergency care ^38^. Existing point-of-care (POC) biosensors - whether optical^39-41^ or electrochemical^42-45^ - address some of these limitations, but still face challenges in balancing sensitivity, speed, cost-efficiency, and quantitative accuracy^46,47^.

To overcome these limitations, we present a **<u>s</u>warm intelligence of <u>m</u>icrobead-inspired, <u>AI</u>-assisted, and <u>s</u>martphone-based** (**SIMAIS**) **biosensing platform** as a novel and effective solution (Supplementary Video 1). The SIMAIS biosensor demonstrates superior speed and sensitivity, achieving results approximately 10 times faster than ELISA with a tenfold increase in sensitivity compared to conventional lateral flow assays (LFAs)^48-50^. Using interferon-gamma (IFN-γ) as a model biomarker, we validated the system across various concentration levels, achieving an overall machine learning classification accuracy of 87%.

## 2. Results

### Overview of the Workflow of the SIMAIS Biosensing Platform

The complete detection workflow of the SIMAIS platform for protein detection is illustrated in **Figure 1c**. The process begins with adding 90 µl test samples and 90 µl control solutions to 10 µl of the pre-functionalised MBs within the vials. Here, the pre-functionalised MBs represent streptavidin-coated MBs with specific antibodies (MB-Strep-Ab) and the preparation protocol of which are provided in the Materials and Methods section. The mixture is gently shaken by hand at room temperature for 3 minutes to facilitate the binding of target antigens to the MB-Strep-Ab, ensuring sufficient interaction. After incubation, the samples are transferred into sampling wells and then placed onto the magnetic plate (ChipMag Plate, preparation protocol provided in the Materials and Methods section) and allowed to stand for an additional 5 minutes. The magnetic field induces the aggregation of the magnetic MBs, facilitating the formation of distinct deposition patterns at the well bottom. These patterns can be analysed both qualitatively and quantitatively (**Figure 1d**).

For qualitative assessment (**Figure 1d**, left), users can visually compare the MB patterns formed in the test and control wells. In the control group, where no antigen is present, the MBs aggregate into a single, tight central ring. In contrast, in the presence of the target antigen, the MBs form two concentric rings: a light inner core and a bigger and denser outer ring. This visually discernible difference in ring morphology enables rapid, naked-eye discrimination between negative and positive samples. These naked-eye-detectable patterns enable a rapid, instrument-free indication of biomarker presence.

For quantitative evaluation (**Figure 1d**, right), a custom smartphone app embedded with a trained machine learning (ML) algorithm is employed. After image capture using the phone’s camera, the app processes the MB pattern and predicts the corresponding analyte concentration range. To ensure standardisation across tests, the app first prompts users to capture an image of a control well. This image is validated by the ML model to confirm correct incubation and imaging conditions before the test image can be analysed. Only after a valid reference is confirmed does the app allow capture of the test sample image. This dual-step validation ensures that the pattern formation and lighting conditions are within acceptable ranges for accurate concentration prediction. The smartphone readout interface then displays the result directly on screen, offering an accessible diagnostic solution without the need for laboratory equipment or trained personnel.

### Understanding the Pattern Formation of Magnetic Microbeads

To systematically investigate this behaviour of MBs, we developed a custom laboratory imaging platform capable of real-time observation and image acquisition of MB motion and pattern formation under an applied magnetic field. This setup comprised a USB microscope mounted on an XY translation stage, a uniform backlight positioned above a 96‐well ChipMag Plate, and sample wells containing antibody‐functionalised MBs with or without the target analyte (**Figure 2a**). This low-cost imaging platform features a motorised XY translation stage controlled by Raspberry Pi, enabling rapid ‘one-click’ image acquisition across multiple wells on a 96-well plate, significantly simplifying the operation and reducing imaging time. Additionally, holders and covers constructed from polymethyl methacrylate (PMMA) are specifically selected to avoid interference with the magnetic field. The use of covers further ensures stable lighting conditions, minimising environmental lighting variability and enhancing data reproducibility. By systematically capturing images under controlled lighting and positioning, the platform captures reproducible data for analysis.

**Figure 2.**
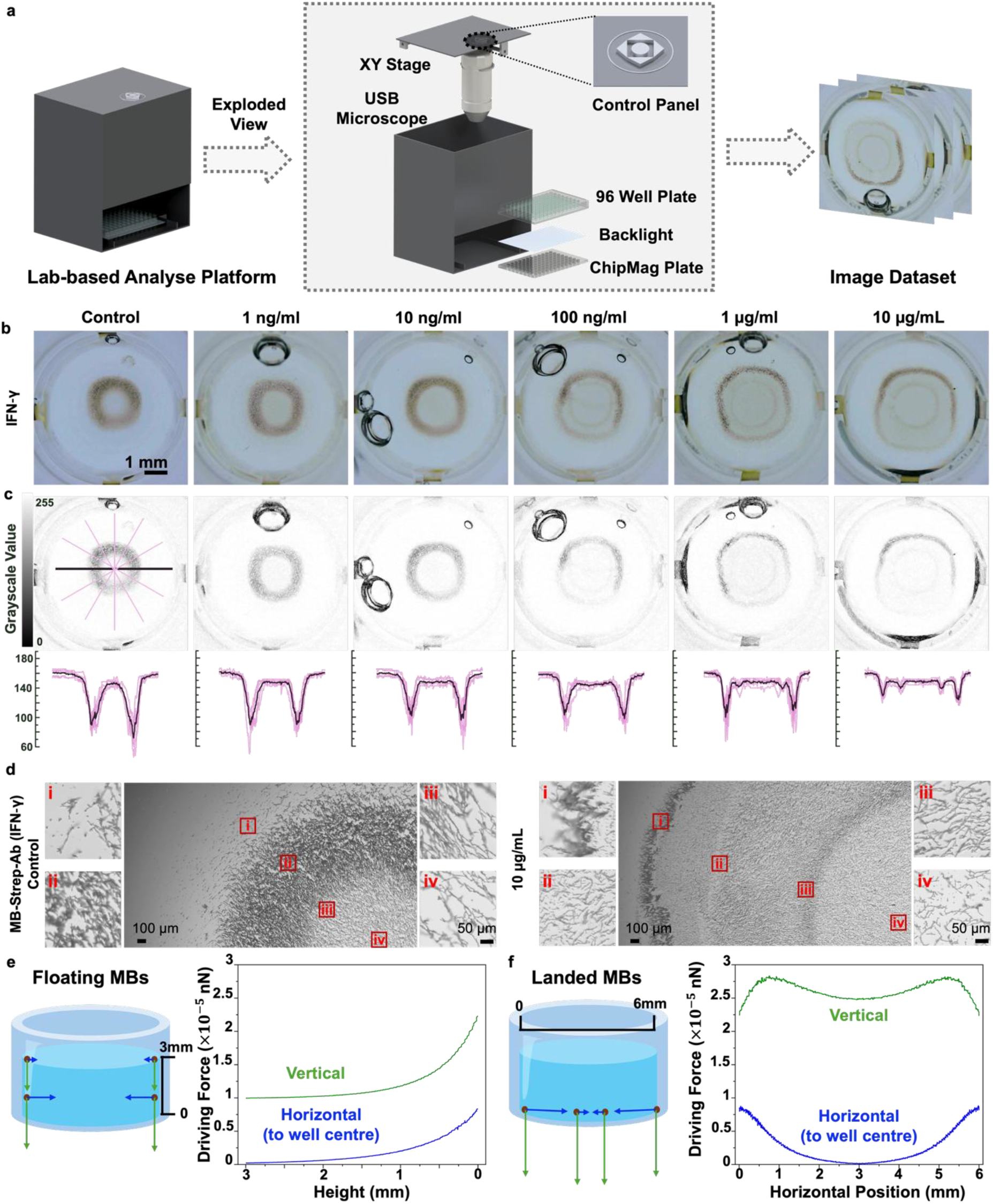
Data collection and mechanism analysis of MBs-formed pattern. a, Illustration of the imaging platform for data collection with a detailed exploded view. b, MB patterns at varying analyte (IFN‐γ) concentrations. c, Grayscale transformations by ImageJ and radial grayscale intensity plots. d, Detailed high-magnification images for the MB pattern. e, COMSOL Multiphysics simulation for floating MBs. f, COMSOL Multiphysics simulation for landed MBs.

Using interferon-gamma (IFN‐γ) as the model protein, we observed that the MBs formed visually distinct ring patterns that correlated with analyte concentrations (**Figure 2b**). Specifically, higher concentrations of IFN‐γ (ranging from 1 ng/mL to 10 µg/mL) produced progressively larger, naked-eye-visible rings, while lower concentrations resulted in smaller and more compact patterns. To quantitatively analyse these patterns, grayscale intensity profiles were extracted using ImageJ (**Figure 2c**). Five equally spaced radial lines (pink coloured lines) were drawn from the centre of each ring, and the grayscale intensity values along each line were measured and averaged. This analysis revealed a concentration-dependent shift in the position of peak intensity, further confirming the strong correlation between antigen presence and the resulting MB pattern morphology.

High magnification images indicate that MBs in the experimental groups exhibit shorter, discontinuous chain-like assemblies compared to the more extended structures observed in the control group (**Figure 2d**). These microstructural changes are indicative of antigen-mediated alterations in interparticle interactions under magnetic confinement.

We further evaluated the surface charge (zeta potential) and hydrodynamic diameter of the MBs following incubation with varying concentrations of the analyte, to investigate how biomolecular binding influences their physicochemical properties and aggregation behaviour (Supplementary Discussion 1 & Supplementary Figure 1). We discovered that the zeta potential remained stable, at −11 to −15 mV across bare MBs, antibody-conjugated MBs, and antigen-bound MBs, whereas the Z-average hydrodynamic diameter and polydispersity index (PDI) apparently increased, which indicates successful bioconjugation. Together, these data show that electrostatic surface charge is not the primary driver of the ring-pattern formation.

We further investigated whether the friction at the bottom of the well contributes to pattern formation. To do this, we compared wells with different surface textures, including those coated and uncoated with bovine serum albumin (BSA), which is known to reduce friction^51^. The results (Supplementary Discussion 2 & Supplementary Figure 2) show that wells treated with BSA exhibit a clearer trend of MB patterns changing with analyte concentration. For the case without BSA treatment, MB tends to settle at the bottom and no longer aggregate toward the centre of the well, thereby preventing the formation of distinct MB patterns. This indicates that increased frictional resistance, significantly altered the morphology of the MB aggregation patterns. These observations suggest that frictional interactions play a key role in shaping the spontaneously organising behaviour of MBs under a magnetic field.

To test the hypothesis, we performed a numerical simulation using COMSOL Multiphysics to analyse the physical forces governing MBs deposition and ring pattern formation. In this simulation, MBs were modelled as superparamagnetic spherical particles suspended in a non-magnetic fluid medium (*χ*_*fluid*_ ≈ 0), ensuring that the magnetic susceptibility *χ* in the force calculations solely represents the MBs’ properties. The primary forces acting on an individual MB include the gravitational force and the magnetic field gradient force, both of which determine its movement and final spatial distribution.

The net vertical force (*F*_*z*_) results from the sum of the downward gravitational force (*F*_*g*_) and the vertical component of the magnetic gradient force (*F*_*magnetic_z*_). Following the approach in previous research^52^, *F*_*magnetic_z*_ on a superparamagnetic sphere in an external field is given by:

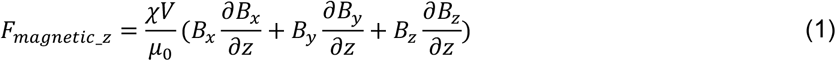

where 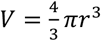 is the volume of the MB, *μ*_0_ is the permeability of free space, and *B*_*x*_, *B*_*y*_, *B*_*z*_ denote the magnetic flux density components in the respective spatial directions. *F*_*g*_ is modelled as *F*_*g*_ = *m*_*bead*_*g* where *m*_*bead*_ is the mass of the MB. The net vertical force, therefore, is:

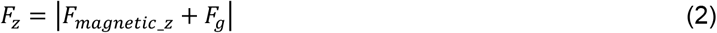

The horizontal force (*F*_*horizontal*_) is entirely dictated by the lateral magnetic field gradient components. Similarly, the net force in the x-y plane is given by:

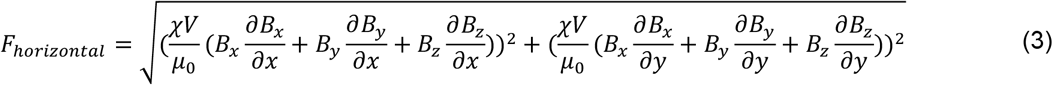

This equation models the MB’s tendency to move toward the centre of the well due to the horizontal magnetic force.

From the simulation results and the experimental observations in **Figure 2e**, it is evident that the vertical force dominates the MB’s behaviour before it reaches the bottom surface of the well, leading to a downward deposition phase. Once in contact with the well bottom (**Figure 2f**), the balance between the horizontal magnetic force toward the centre and resistance from movement governs the final bead arrangement. The presence of antigen in the sample accelerates the equilibrium process, leading to a broader ring-like pattern formation compared to the tighter pattern in the control.

A real‐time experimental demonstration of MB pattern formation is provided in Supplementary Video 2, along with a schematic animation in the Supplementary Video 3. These resources visually confirm that the interplay of vertical and horizontal forces drives the distinct aggregation patterns, reinforcing the mechanism insights obtained from computational simulations and laboratory imaging.

In summary, our combined imaging, zeta-potential/dynamic light scattering (DLS) characterisation, surface-roughness experiments and COMSOL Multiphysics simulation show that the MB pattern formation is likely to be governed by a three-way balance of magnetic-field gradients, gravity-driven sedimentation and frictional interactions at the well floor. Antibody conjugation and antigen binding leave the zeta-potential virtually unchanged, making it unlikely that electrostatic forces dictate the pattern. Instead, once beads reach the substrate, lateral magnetic attraction pulls them centre-ward while surface friction – minimised on smooth, BSA-passivated wells and intensified on uncoated ones – opposes this motion, fixing the final ring-like pattern. This interplay of magnetic, gravitational and interfacial forces, modulated by biomolecular binding, underpins the SIMAIS biosensor’s concentration-encoded, swarm-like self-assembly.

### Development of the SIMAIS Biosensing Platform

After understanding the physical and biochemical mechanisms underlying the formation of MB patterns and establishing a clear correlation between ring morphologies and analyte concentration, we explored whether this visually interpretable, concentration-dependent phenomenon could be harnessed for protein detection. While the naked-eye-visible ring patterns already offer a rapid and accessible qualitative readout, quantitative interpretation, especially at low concentrations or in clinical samples with subtle morphological differences, requires greater analytical precision and reproducibility than visual inspection alone can provide.

To address this, we introduced a machine learning (ML) framework to enhance the platform’s analytical precision and objectivity. ML can detect and quantify subtle, high-dimensional features within MB patterns (e.g., ring width, granularity, texture distribution) that may be imperceptible to the human eye or difficult to measure using traditional image processing methods.

#### (1) ML Classifier Structure and Performance

A YOLO (You Only Look Once) network is employed to perform image classification for IFN-γ concentration detection. YOLO is a state-of-the-art, real-time network that has revolutionised the field due to its ability to achieve high accuracy and speed simultaneously^53^. The architecture of YOLO for classification, shown in **Figure 3a**, consists of a feature extractor as its backbone and a classification head for making class predictions based on the input image features. A detailed block diagram of the network can be found in Supplementary Discussion 3 & Supplementary Figure 3.

**Figure 3.**
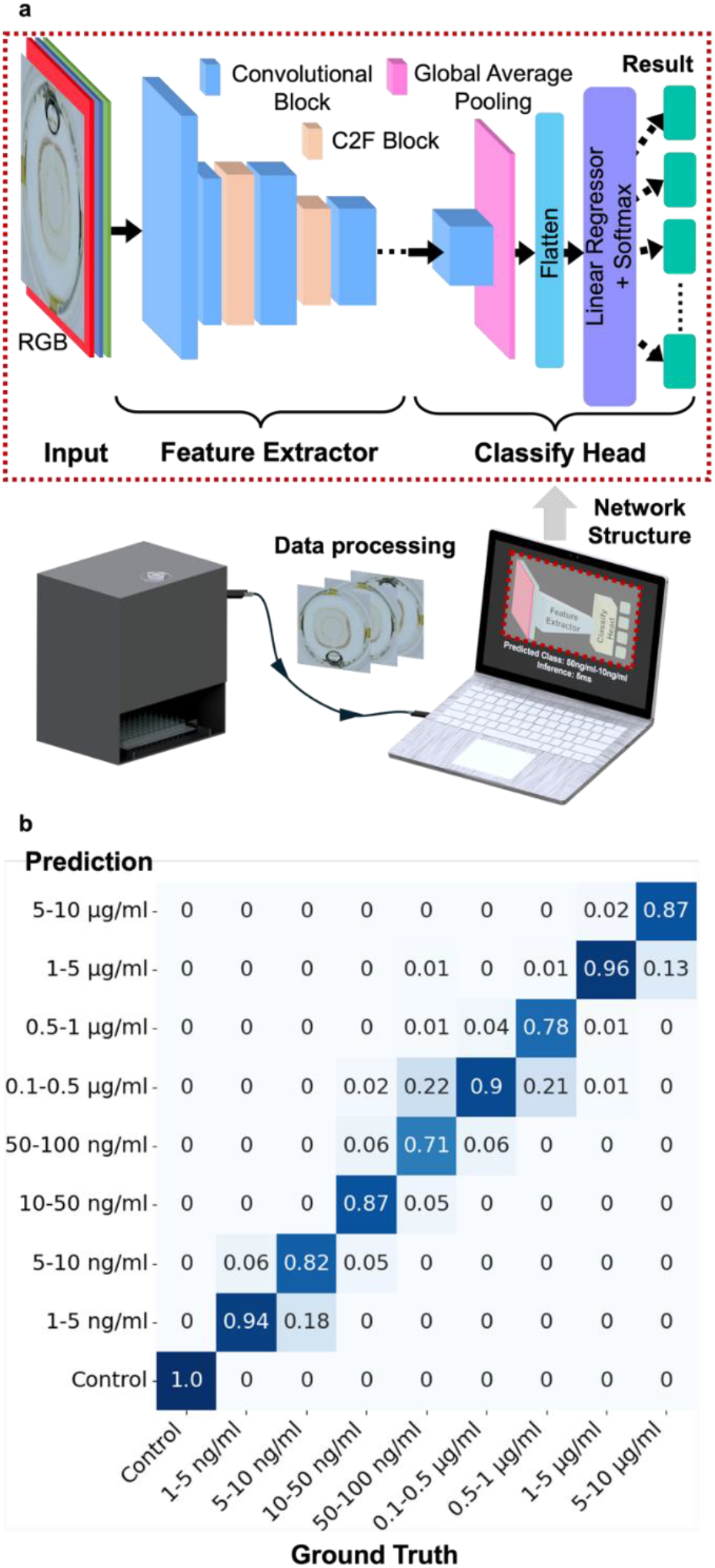
ML network structure and performance. a, Schematic representation of data processing and model training workflow with YOLO-based classification network structure. b, Confusion matrix of the trained model performance.

To train the YOLO network, we prepared a dataset consisting of approximately 40 raw images for each level of IFN-γ concentration to ensure class balance. Several augmentation techniques, including flipping (horizontally and vertically), rotation (−30° to +30°), brightness and contrast adjustment (−20% to +20%), and blurring (up to 3 pixels), are applied to enhance the dataset and prevent overfitting. These augmentations increase the dataset size by five times, resulting in a more diverse and robust dataset. Based on varying classes, the dataset is then split into 70% for training and 15% for the validation. The rest of the 15% image data is used as the testing set.

We evaluated the performance of the trained network using a confusion matrix (**Figure 3b)**. Each row of the matrix represents the actual class label for the image, while each column represents the predicted class. The diagonal elements of the matrix indicate correct classifications, where the predicted class matches the actual class, representing the accuracy for each class. In contrast, the off-diagonal elements represent misclassifications. When classes are split as illustrated in the figure, each concentration level achieves > 70% prediction accuracy. With classes split based on the logarithm (base 10) of concentrations to compensate misclassifications occurring to neighbouring concentration level, the model performs even better. This performance is primarily limited by the dataset size. The trained network achieves an average accuracy (∑ Accuracy for each class /n (Total number of classes), where n = 9) of 87.2%, indicating that the model performs well across the entire range of concentrations.

#### (2) Smartphone App and POC Device

The development of the SIMAIS device and its accompanying smartphone app is an important step toward enabling rapid and user-friendly protein detection. The overall working pipeline of the smartphone app is depicted in **Figure 4a**.

**Figure 4.**
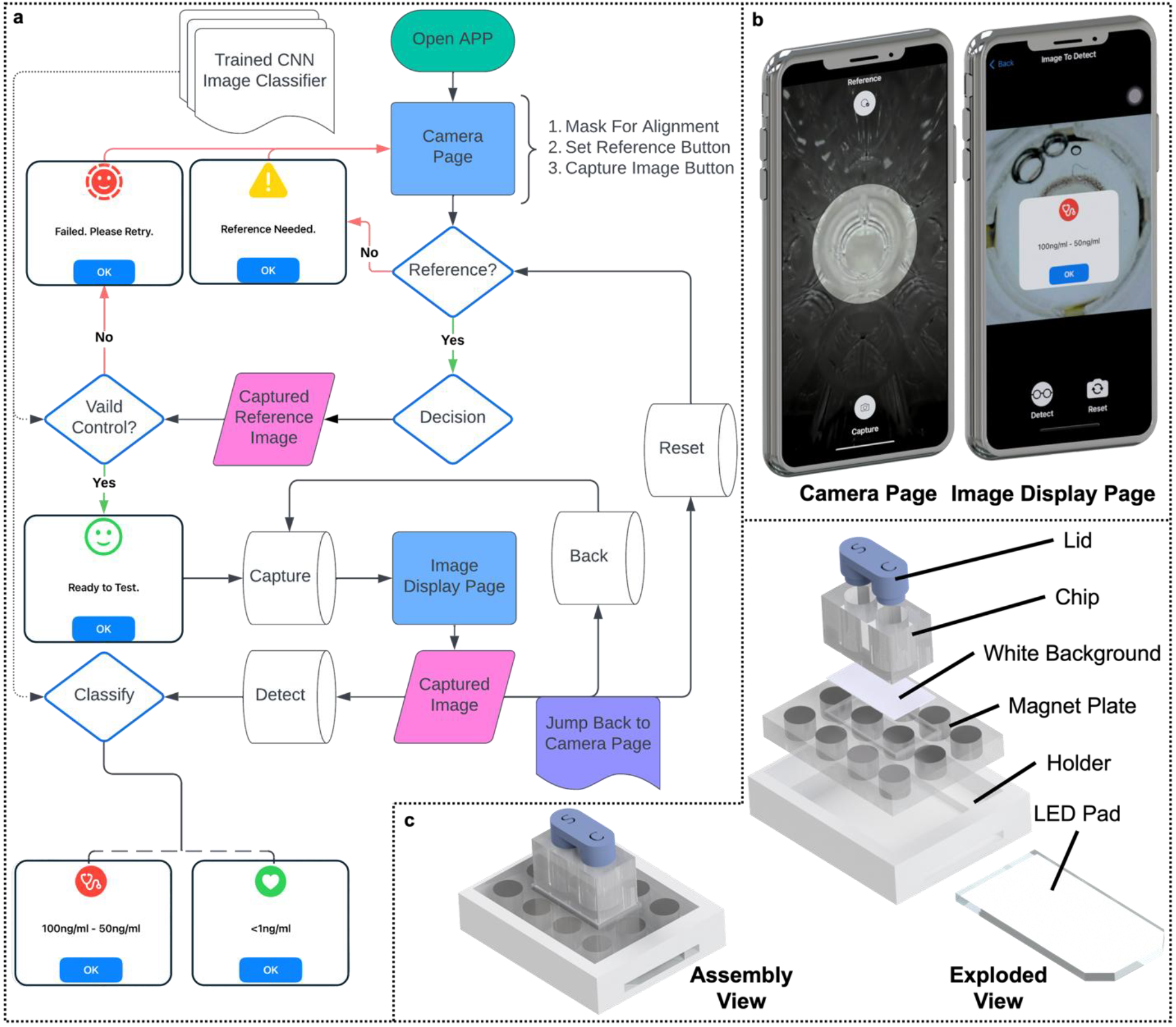
Illustration of the smartphone app and POCT device. a, Workflow diagram of the smartphone app. b, Screenshots of the app interfaces. c, Schematic illustration of the POC device components.

Upon launching the app, the camera page, which has a central circular mask designed to align with the outer-diameter of 96-well plate wells or the POC wells during image capture, is presented. This alignment is essential for capturing consistent images feeding into the analysis running in the background. The interface on this page includes two buttons: the ‘*Reference*’ button located at the top and the ‘*Capture*’ button at the bottom of the screen, as shown in **Figure 4b**. The ‘*Reference*’ button allows users to capture the image of the control sample, which serves as a reference image. A valid reference image will be identified as ‘*control*’ by the deployed ML classifier. The ‘*Capture*’ button is activated only after capturing a valid reference image. This ensures that the previous sample processing, such as shaking time, is appropriate for analysis. After capturing the image of the sample, the app transitions to the image display page (**Figure 4b**). This page displays the recently captured image and provides three functional buttons for further interaction. The ‘*Detect*’ button, located at the bottom left, initiates the concentration prediction by feeding the image to the in-app ML network and shows the prediction result. The ‘*Back*’ button at the top left is for returning to the camera page to retake the test sample image without resetting the reference image, which is useful if the initial capture was misaligned. The ‘*Reset*’ button at the bottom right clears the current reference image, allowing users to begin a new test with a different POC chip.

The POC device set, as illustrated in **Figure 4c**, comprises several components designed for ease of use and optimal functionality. The main component is the POC chip with a lid, which includes a sampling well where the MB working buffer interacts with the test samples and a control well for setting a reference. The lid serves to protect the samples from contamination and spills during the shaking process. The MagChip plate, which is a magnet plate designed with a white background, and a slot specifically for accommodating the chip, facilitates the MBs aggregation process and enhances image contrast during capture. A holder tightly houses the MagChip and aligns it with the LED pad, which provides uniform illumination from beneath for capturing high-quality images of the MB patterns formed.

The design of both the smartphone app and the POC device emphasises user-friendliness, reusability and price-friendliness, making the system accessible to more users with minimal training. The integration of the ML network within the app enhances diagnostic capability by providing precise quantitative analysis without the need for complex laboratory equipment and a long waiting period. A demonstration of how to use the POC device with the smartphone app is captured in Supplementary Video 4.

### Evaluate the Performance of the SIMAIS Biosensing Platform

To ascertain the practical diagnostic utility of SIMAIS biosensing platform, we examined four core performance attributes – sensitivity, working range, specificity/selectivity, and matrix tolerance – based on MB patterns recorded in the experiments.

#### (1) Sensitivity and working range

With a categorical, ML classifier read-out, the classical 3σ/10σ definitions^54^ of LoD (Limit of Detection)/LoQ (Limit of Quantification) are not directly applicable. Instead, we define sensitivity as the lowest concentration class that can be distinguished from blanks with high recall. In the confusion matrix (**Figure 3b**), the 1-5 ng/mL level shows a classification recall of 0.94, without being misclassified to ‘Control’. We, therefore, report a practical LoD of 1-5 ng/mL for IFN-γ under the present model classification scheme. Correct-class recall remains larger than 0.7 across every higher concentration level, extending to the largest class of 5-10 µg/mL. Taken together, SIMAIS biosensing platform covers almost six orders of magnitude. The underlying MB pattern versus log-concentration relationship is monotonic but not strictly linear. Similarly, as the output is categorical rather than continuous, we term this working range as a dynamic classification range.

#### (2) Specificity/selectivity

**Figure 5a** demonstrates stringent immune recognition: MBs coated with polyclonal IFN-γ antibodies return the correct concentration label for every IFN-γ sample across the dynamic classification range while assigning all TNF-α antigen challenges, even at 10 µg/mL, to the ‘*Control*’ class, indicating negligible cross-reactivity.

**Figure 5.**
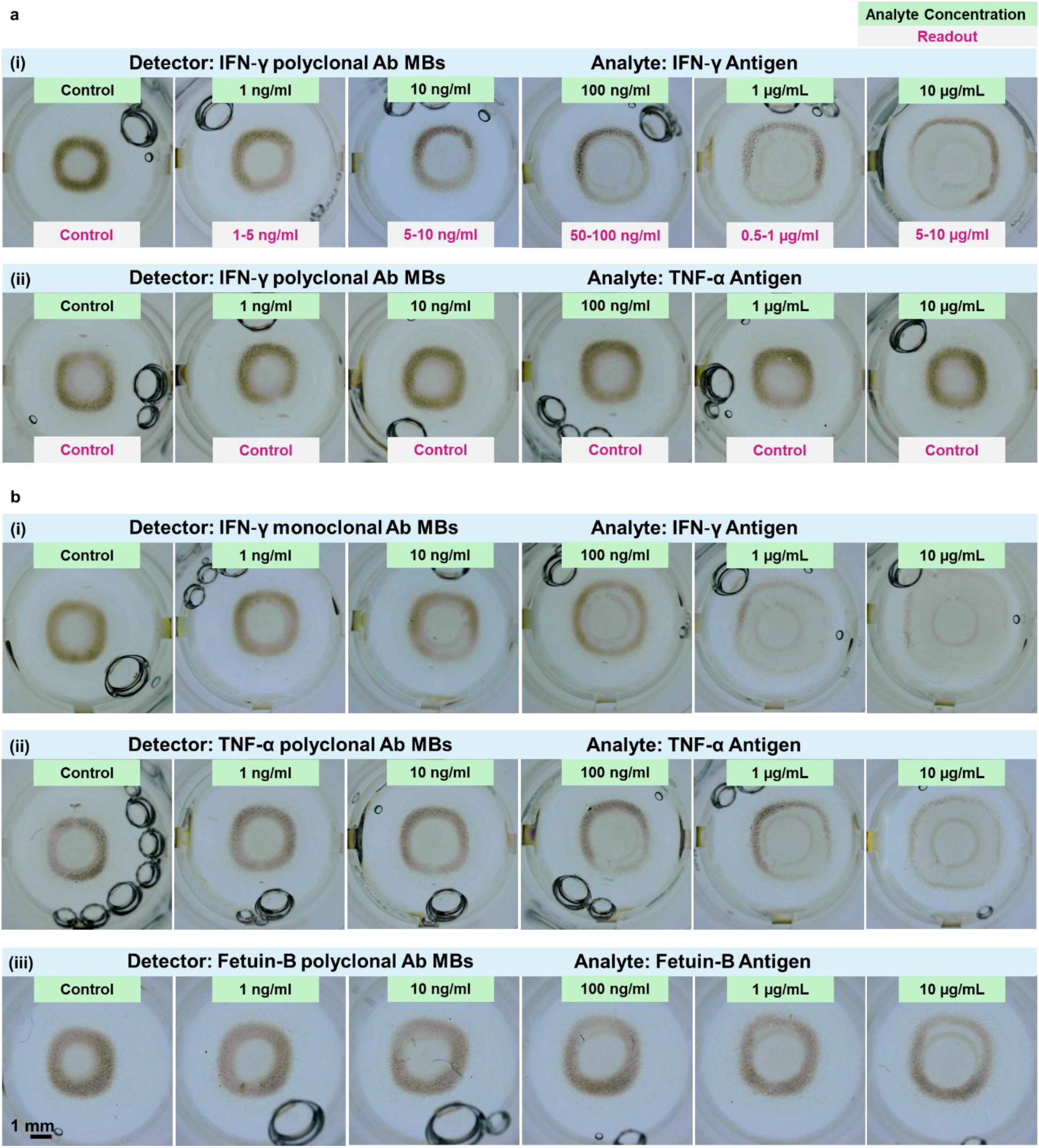
SIMAIS platform performance evaluation. a, Specificity/selectivity. MBs functionalised with IFN-γ polyclonal antibodies are tested with serial dilutions of either on-target IFN-γ antigen (i) or off-target TNF-α antigen (ii). The ML classifier correctly assigns each on-target sample to its concentration level, while labelling every off-target well as “Control”, confirming negligible cross-reactivity. b, Transferability to other antibody-antigen pairs. MB pattern trends are reproduced with alternative detectors: (i) IFN-γ mAb-MBs, (ii) TNF-α pAb-MBs, and (iii) Fetuin-B pAb-MBs, which demonstrate the transferability of the SIMAIS biosensing method.

#### (3) Transferability to other antibody-antigen pairs

The same MB pattern formation trend is observed when either the antibody clonality or the target protein is changed (**Figure 5b**). Each detector – whether a monoclonal IFN-γ MB detector, a polyclonal TNF-α MB detector, or a Fetuin-B MB detector – produces concentric patterns that expand monotonically across the dynamic classification range. These experiments confirm that the SIMAIS biosensing platform is a generally deployable pattern-coding platform.

#### (4) Matrix tolerance

Sample matrix effects were investigated by spiking in IFN-γ to serum, urine and standard 1% BSA in PBS buffer (Supplementary Figure 4 & Supplementary Discussion 4). Compared to the standard buffer, the higher-protein and higher-salt matrices slightly altered the edge of the MB pattern, however a clear visual distinction between antigen-positive samples and their matrix-matched controls is retained in every case. The preservation of this qualitative ‘positive-versus-blank’ contrast indicates that the MB-based biosensing signal is robust to the biochemical complexity of clinically relevant fluids. Thus, it is promising to be optimised further for practical application.

## 3. Discussion

This study presents a novel approach to biosensing, inspired by swarm intelligence. Leveraging this phenomenon, we developed the SIMAIS biosensor, which employs MBs coated with specific antibodies that dynamically organise into distinct patterns when exposed to target biomarkers under an applied magnetic field. This innovative approach translates subtle molecular interactions into macroscopic, visible patterns, offering intuitive qualitative and precise quantitative analysis.

The simulation and experimental validations presented confirm robust, reproducible pattern formation driven by gravitational and magnetic field gradient forces, offering mechanistic insights into the biosensor’s functionality.

Our biosensor demonstrates significant advancements compared to existing diagnostic methodologies. It notably achieves a detection speed ten times faster than traditional ELISA, while simultaneously delivering a tenfold increase in sensitivity relative to conventional lateral flow assays. Additionally, its compatibility with smartphone-based AI further enhances the system’s accessibility and usability, providing rapid, real-time analysis without requiring extensive training or bulky laboratory equipment.

Importantly, the platform is modular: by simply modifying the antibodies coated on the MBs, the SIMAIS biosensor can be readily adapted for detecting a wide range of protein targets. This innovative integration of swarm intelligence and AI technology offers a transformative advance in diagnostic science, paving the way for rapid, cost-effective, and decentralised healthcare diagnostics and real-time patient monitoring.

Distinct features set this biosensing platform apart from other current diagnostic tools. Firstly, biomarker-driven emergence of antibody-functionalised MBs into clearly discernible patterns enables simple visual interpretation, serving as an immediate qualitative indication of biomarker presence. Secondly, quantitative analysis facilitated by a smartphone app equipped with a ML classifier provides accurate biomarker concentration level detection, significantly democratizing access to high-performance diagnostics. Thirdly, the adaptability of this platform is demonstrated through successful detection of multiple biomarkers, including IFN-γ, TNF-α and Fetuin-B, by altering antibody conjugation on the MBs, underscoring the biosensor’s versatility. Furthermore, this antibody-functionalized MBs-pattern biosensing approach detects biomarkers in spiked samples (1% serum, 10% serum, and pure urine), indicating promising robustness for practical diagnostic applications. The current trained ML models can rapidly analyse hundreds of samples in seconds, making them ideal for point-of-care or field applications where speed and scalability are essential. Additionally, by increasing the background complexity and quantity of the dataset in future studies, trained models can adapt to variations in sample background, lighting conditions, and biological matrices (e.g., serum, urine), helping maintain robust performance across diverse clinical and environmental settings. Furthermore, ML algorithms can be embedded into smartphone apps or portable devices, enabling on-device, offline analysis without requiring external servers or internet access.

In summary, the SIMAIS biosensing platform combines LFA-level sensitivity (practical LoD ≈ 1-5 ng/mL) with an exceptionally broad usable span (from 1 ng/mL to 10 µg/mL), preserves strict antibody-mediated specificity, adapts seamlessly to multiple antibody-antigen pairs, and retains a clear positive-versus-blank signature even in complex matrices such as diluted serum and pure urine. These attributes, delivered through a rapid 10-minute, mix-and-measure workflow and an automated vision-based ML classifier embedded in a smartphone, position SIMAIS biosensing as a versatile and robust candidate for point-of-care protein diagnostics. Acknowledging the current limitation in classifier accuracy due to the small dataset, this performance can be enhanced substantially with future additional high-quality training data, positioning this technology to have transformative implications in healthcare diagnostics., particularly suitable for resource-limited and decentralised healthcare settings.

## 4. Materials and Methods

### Biochemicals list

IFN-γ polyclonal antibody, biotin (500-P32BT, PeproTech®). IFN-γ monoclonal antibody, biotin (13-7319-81, eBioscience™), Human IFN-γ recombinant protein (300-02, PeproTech®). TNF-α polyclonal antibody, biotin (500-P31ABT, PeproTech®). Human TNF-α recombinant protein (300-01A, PeproTech®). Fetuin-B polyclonal antibody (BAF1725, Bio-Techne®). Human Fetuin-B protein (1725-PI, Bio-Techne®). Bovine serum albumin (05470, Sigma-Aldrich®). Normal Human Serum (Invitrogen, Thermo Scientific™). Phosphate-buffered saline (P4417, Sigma-Aldrich®).

### Preparation of MB-Strep-Ab

Working buffer contains (i), streptavidin coated magnetic microbeads (MB-Strep) (Dynabeads™ MyOne™ Streptavidin C1, selected as discussed in Supplementary Discussion 5 & Supplementary Figure 5) and (ii), biotinylated antibody. The MB-Strep are first washed three times with sterile DI water and then resuspended in 1% (w/v) bovine serum albumin (BSA) in phosphate-buffered saline (PBS). The resuspended MB-Strep and biotinylated antibody are mixed at a mass ratio of 80:1 (selected as discussed in Supplementary Discussion 6 & Supplementary Figure 6) in a microcentrifuge tube. The mixture is incubated at room temperature for 5 minutes with gentle rotation at 30 rpm using a tube rotator (Thermo Scientific™) to facilitate binding. Finally, MB-Strep-Ab is resuspended in 0.5 mg/mL with 1% BSA in PBS to complete the preparation of the MB-Strep-Ab conjugate working buffer.

### Preparation of antigen-spiked samples

Human Recombinant IFN-γ, TNF-α or Fetuin-B stocks (100 µg/mL in 1% BSA in PBS) are serially diluted in the same buffer to obtain working standards from 10 µg/mL down to 1 ng/mL. For matrix-tolerance experiments, human serum is thawed on ice, equilibrated to room temperature, centrifuged at 10,000 rpm for 5 min and the supernatant dilutes to 10% or 1% (v/v) in 5 % BSA in PBS. Urine is simply centrifuged at 10,000 rpm for 5 min to remove particulates. Each matrix is then spiked 1:9 (v/v) with the appropriate antigen working solution to give final concentrations identical to the buffer standards while preserving the desired matrix fraction (10%, 1% or pure urine). Un-spiked matrix processed in parallel serves as the ‘Control’ condition.

### MagChip fabrication

Two types of MagChips are prepared. The first is a MagChip plate designed for 96-well plates to enable high-throughput image data collection. A magnet holder was 3D-printed using tough polylactic acid (PLA) filament (Ultimaker 3D printer) and crafted to accommodate 96 N52 neodymium disk magnets (6 mm × 4 mm, EP657N52, e-Magnets UK). Each magnet is inserted into the holder with the same north pole orientation to ensure a non-distorted magnetic field across the plate. The second MagChip is fabricated following the same protocol as the 96-magnet plate but was designed to hold only 12 magnets. The corresponding magnetic field simulation via COMSOL Multiphysics is illustrated in Supplementary Discussion 7 & Supplementary Figure 7. This smaller-scale MagChip offers a more compact alternative for POC settings.

### Lab-based analyse platform fabrication

The lab-based analysis platform is assembled to facilitate the high-throughput imaging in 96-well plate during the data collection stage. It consists of an XY stage, a portable USB microscope, an LED backlight pad, and a control system. The XY stage is created using two stepper motor linear actuators with sliders that provide a resolution of 0.025 mm per step, allowing precise positioning of samples. A portable USB microscope (Jiusion Digital Microscope), providing 40× to 1000× magnification, is mounted above the stage to provide imaging capabilities. To ensure uniform illumination of the samples, an LED backlight pad was placed beneath the sample area. A Raspberry Pi 4B is employed as the microcontroller for motor control, image capture, and communication with a computer.

### POC chips fabrication

For portability and affordability, we used PMMA to manufacture the POC chip and 3D printed its cover lid using PLA filament. The chip has an overall volume of 20 mm × 10 mm × 12 mm (length × width × height), with a spacing of 9 mm between the two wells. The well diameter is 6.35 mm, and the depth is 10.67 mm (same as the dimensions of a single well in a 96-well plate).

## Supporting information

Supplementary Discussions 1-7 Supplementary Figures S1-S7

## Acknowledgments

C.Z. acknowledges the funding support from The Royal Society, UK (Grant Nos. RG/R1/241228 and IEC/NSFC/233339), Wessex Medical Innovation Fund 2025 (AF06) from Wessex Medical Trust, UK, and internal fundings from the School of Electronics and Computer Science, University of Southampton, UK. H.F.S acknowledges grant funding from the Academy of Medical Sciences (SGL031/1084) and the support of the Southampton Biomedical Research Centre. S.-Y.T. gratefully acknowledges the research funded by the Australian Research Council Future Fellowship (FT230100257), Discovery Project (DP250103029) and UNSW Scientia Fellowship.

## Author Contributions

C.Z. conducted the initial experiments and obtained the preliminary data. The conceptual framework for this work was developed by C.Z., S.-Y.T., and J.Z.; J.Z. and B. performed the experiments, analysed the data, and prepared the figures with support from X.W., T.W., X.L., and Y. Z.; H.F.S. and Y.W. provided essential resources and contributed to the investigation and discussion. The first draft of the manuscript was written by J.Z. and revised with input from all authors.

## Competing Interest Statement

The authors declare no competing financial interests or personal relationships that could have appeared to influence the work reported in this paper.

## Notes

### Competing Interest Statement

The authors have declared no competing interest.

