## Supplementary Discussions 1-7 Supplementary Figures S1-S7 for "SIMAIS: A swarm intelligence inspired biosensor for rapid protein detection"

**This file includes:**
Supplementary Discussions 1-7
Supplementary Figures S1-S7
Caption for Supplementary Videos 1-4

### Supplementary Discussion 1: Zeta potential and hydrodynamic diameter test

To evaluate the stepwise modification of MBs based detection complexes, three sample groups are prepared: (1) bare MBs, (2) MB conjugated with antibodies (MB-Abs), and (3) MB-Abs further incubated with target antigen (MB–Abs–Ag). Result is shown in **Supplementary Figure 1**. The samples are tested in standard buffer (1% BSA in PBS). The hydrodynamic diameter (Z-average), polydispersity index (PDI), and zeta potential of each sample were measured using a Malvern Zetasizer Nano ZS instrument. All measurements are performed at 25°C with triplicate repeats (n=3) to ensure reproducibility.

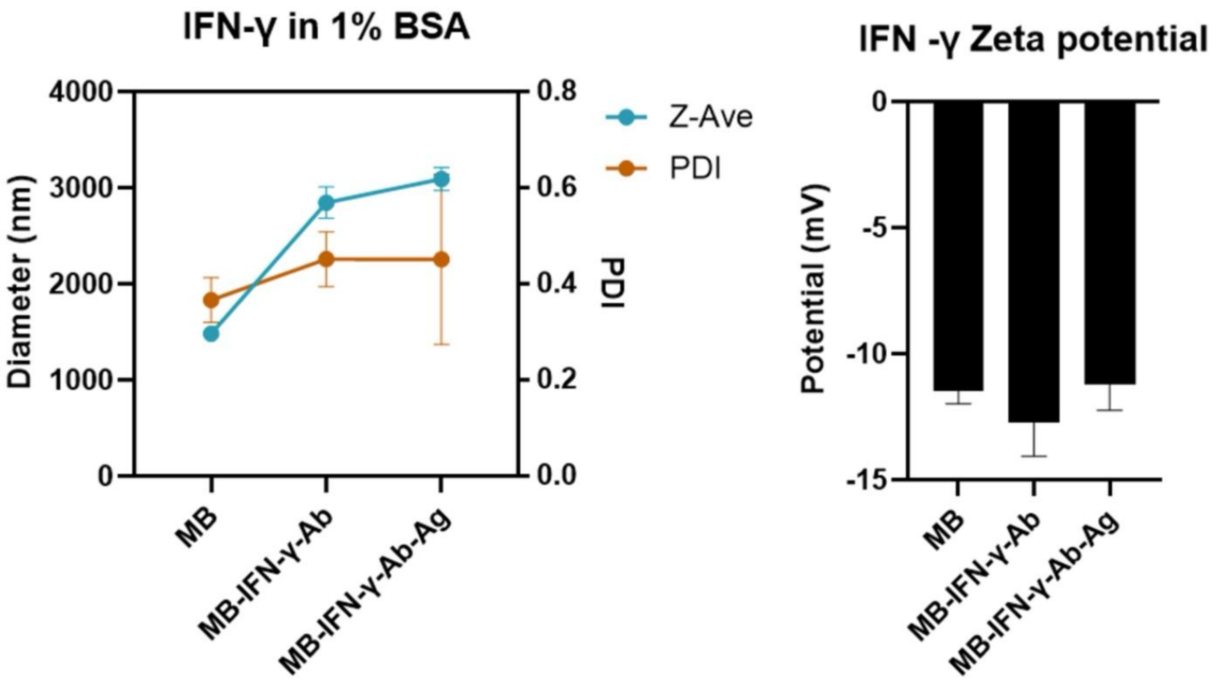


**Supplementary Figure 1.** Characterisation of IFN-γ binding on MBs via DLS and zeta potential analysis. Dynamic light scattering (DLS, left) and zeta potential (right) measurements of MB, antibody-conjugated MBs (MB–IFN-γ–Ab), and antigen-bound complexes (MB–IFN-γ–Ab–Ag) in 1% BSA in PBS solution. Z-average diameter (Z-Ave, blue) and polydispersity index (PDI, orange) are used to assess size distribution. Zeta potential (black bars) indicates surface charge variation across modification steps.

### Supplementary Discussion 2: Influence of well-bottom friction on MB pattern formation

Passivating the well bottom with 1% BSA in PBS markedly enhances the clarity and concentration-dependence of the MB patterns (**Supplementary Figure 2**, top-row). By reducing surface roughness and frictional drag, BSA allows the lateral magnetic force to pull bead clusters towards the centre, yielding well-defined pattern scaled with antigen load. In native wells, the higher friction opposes this lateral motion; MBs settle where they land and only faint, irregular aggregates are observed (**Supplementary Figure 2**, bottom-row). These observations confirm that interfacial friction is a critical parameter governing the final geometry of the MB pattern.

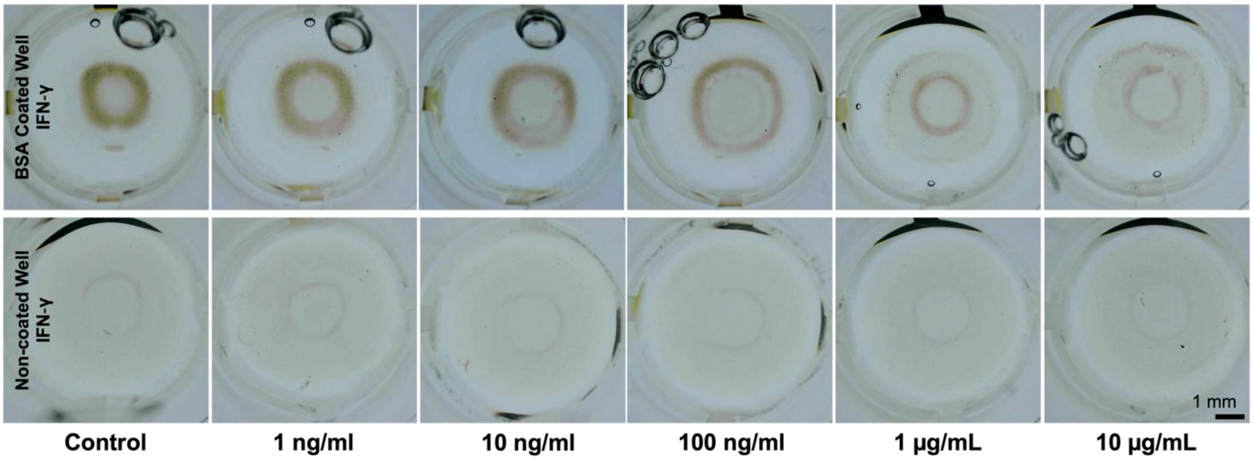


**Supplementary Figure 2.** Representative images of MB patterns formed in BSA-coated wells (top-row) versus non-coated wells (bottom-row). Columns correspond to increasing IFN-γ concentrations. In the low-friction, BSA-coated wells, concentric rings enlarge monotonically with antigen concentration, whereas in the higher-friction native wells the MBs largely sediment without organising into distinct rings.

### Supplementary Discussion 3: YOLO Network

**Supplementary Figure 3** illustrates the architecture of the YOLOv8 network, highlighting the key components: the feature extractor and the classify head. The network incorporates several types of blocks, such as convolutional blocks, bottleneck blocks, and C2f blocks, as explained in the figure. These blocks work together to process the input images and generate accurate predictions of cytokine concentrations.

The feature extractor processes the input images to extract salient features that are indicative of different analyte concentrations. It employs a series of convolutional blocks and C2f blocks to capture visual patterns. The classify head, as the follow-up, maps the complex features to specific output classes corresponding to various concentration levels based on the features extracted. The three important blocks for building the YOLO network are also illustrated in **Supplementary Figure 3**. The convolutional block, as the fundamental building block of the network, consists of a convolutional layer followed by an activation function (SiLU in this case). This block detects local features in the input images, such as edges, textures, and shapes formed by the MB. Bottleneck block is largely used in C2f block, which fuses features from earlier and later layers, allowing the network to consider both shallow features and deep features simultaneously.

With the trained YOLO network, our SIMAIS biosensor allows near-instantaneous analysis of MB patterns. In addition, the modular structure of YOLO facilitates adjustment and expansion. By expanding or updating the training dataset and modifying the output categories, if necessary, the network can be fine-tuned or retrained to further improve prediction accuracy with a larger dataset on the same target protein or to detect other types of target protein.

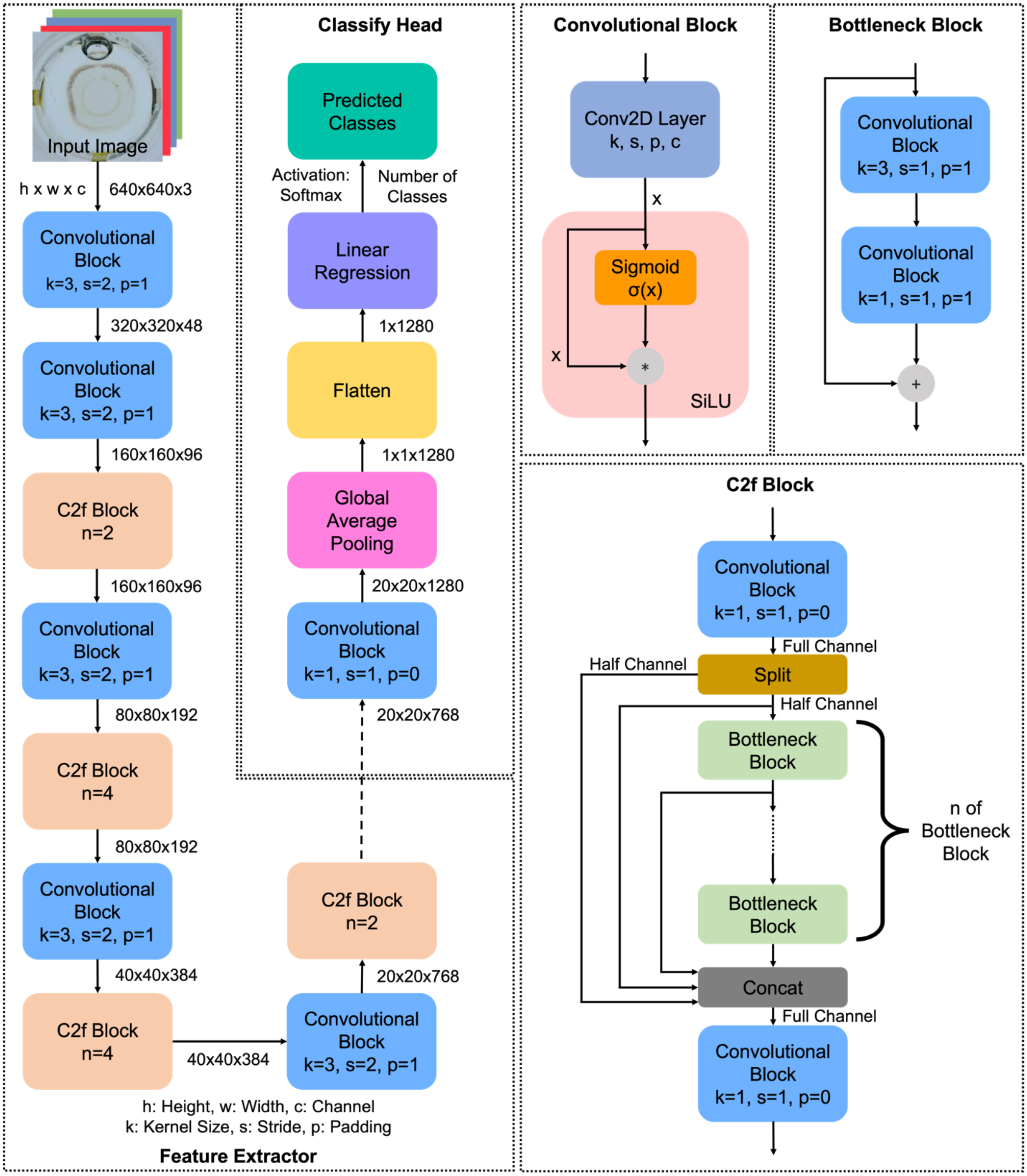


**Supplementary Figure 3.** Schematic representation of the YOLOv8 network used for MB pattern analysis. The diagram illustrates the flow from input RGB image (640×640×3) through the feature extractor - comprising convolutional, bottleneck, and C2f blocks - to the classify head that outputs the cytokine concentration level prediction.

### Supplementary Discussion 4: Matrix-tolerance of the SIMAIS biosensing platform

In samples with high serum content, the final pattern rings formed by the magnetic beads (MBs) were consistently smaller than those observed under standard conditions (1% BSA in PBS). This size reduction adversely affected the performance of our detection algorithm. To address this issue, we found that adjusting the concentrations of both Tween 20 and BSA could gradually mitigated the serum's influence: pattern rings in serum-containing samples became noticeably larger when these additives were present (data not provided). These results suggest the potential for formulating an optimised buffer to reduce serum interference, warranting further investigation.

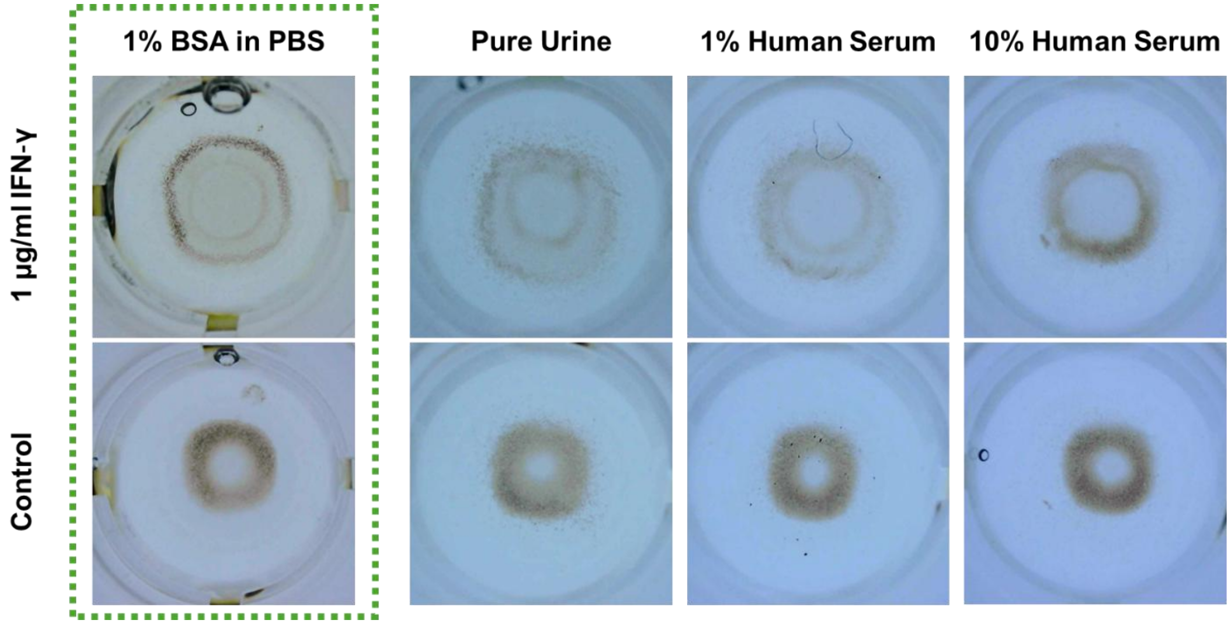


**Supplementary Figure 4.** Matrix-tolerance evaluation. Representative MB pattern images obtained with 1 μg/mL IFN-γ (top-row) and matrix-matched 'Control' samples (bottom-row) in, from left to right: 10% human serum, 1% human serum, pure urine and the standard 1% BSA in PBS buffer.

### Supplementary Discussion 5: MBs selection

We investigate to compare three types of MB-strep to determine the most suitable for our assay: Nanocs Inc. M100™ Magnetic Nanobeads Streptavidin (100 nm), Dynabeads™ MyOne™ Streptavidin C1 (1 µm) and Dynabeads™ M-270 Streptavidin (2.8 µm). This comparison aims to assess how bead size affects pattern formation and assay sensitivity. As illustrated in **Supplementary Figure 5**, at the same bead concentrations, all three types of MB-strep-Ab exhibit a similar trend in pattern formation relative to antigen concentration. Comparing to the 2.8 µm MB-strep-Ab (**Supplementary Figure 5a**), the 100 nm (**Supplementary Figure 5b**) and 1 µm ones (**Supplementary Figure 5c**) demonstrate greater variability in pattern morphology across different antigen concentrations. We speculate that the larger size of the beads results in increased momentum (i.e. mass and speed) of the outer beads as they migrate toward the centre under the influence of the magnetic field. This higher momentum may lead to the compression of the central MB network's pattern, thereby diminishing the system's sensitivity to antigen concentration changes. Although the 100 nm MB-strep present a good visibility for antigen detection, as the size of individual MB-strep is in nanoscale, it takes longer for the pattern formation and possibly need to have a larger bead concentration for making the pattern easier to be distinguished with bare eyes. From this aspect, using 100 nm MB-strep, though could possibly increase the sensitivity of the biosensor, the cost, both on time and material, would be higher. Based on these observations, we select the 1 µm Dynabeads™ MyOne™ Streptavidin C1 for subsequent experiments. The influence of MB size on the assay's sensitivity suggests a potential avenue for further research. Optimising the MB size could enhance the detection capabilities of the system, potentially leading to improved performance in biomarker quantification.

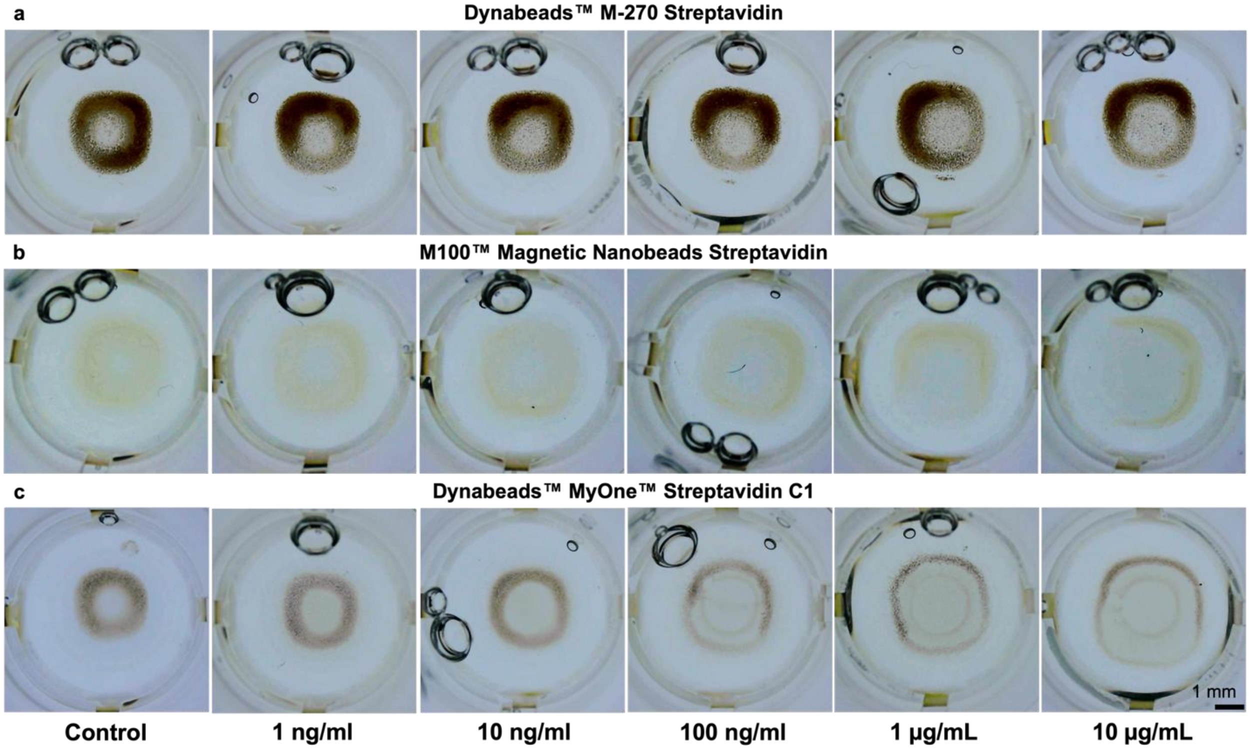

**Supplementary Figure 5.** Investigation of MB-strep. Comparison of the pattern formed by MB of different sizes with the same beads' concentration (a, Dynabeads™ M-270 Streptavidin, 2.8 μm. b, Nanocs Inc. M100™ Magnetic Nanobeads Streptavidin, 100 nm. c, Dynabeads™ MyOne™ Streptavidin C1, 1 μm).

### Supplementary Discussion 6: Antibody-MB conjugation optimisation

Using the selected 1 μm MB-strep, we explore the optimal mass ratio for incubating the MB-strep with biotinylated IFN-γ antibody (Ab). Following standardised procedures for MBs of equal mass, we prepare MB-strep-Ab conjugates with varying MB:Ab mass ratios of 200:1, 160:1, 125:1, 80:1, and 50:1. A control sample consisting of MB without Ab conjugation is also prepared. Each MB-strep-Ab preparation and the control sample are added to a 96-well plate in duplicate. To label the Ab bound to the MB, we add an appropriate amount of fluorescent secondary antibody (Goat anti-Rabbit IgG (H+L) Cross-Adsorbed Secondary Antibody, Alexa Fluor™ 488, Invitrogen™) to each well. The samples are incubated with shaking to facilitate binding. After incubation, excess fluorescent secondary antibody is removed through washing steps. The fluorescence intensity (FI) of each sample is measured using a microplate reader (CLARIOstar® Plus). The FI readings are presented in **Supplementary Figure 6**. These measurements allow us to quantify the amount of Ab successfully conjugated to the MBs at each incubation ratio. By analysing the FI data, we identify the optimal incubation mass ratio as 80:1 which provides sufficient antibody conjugation.

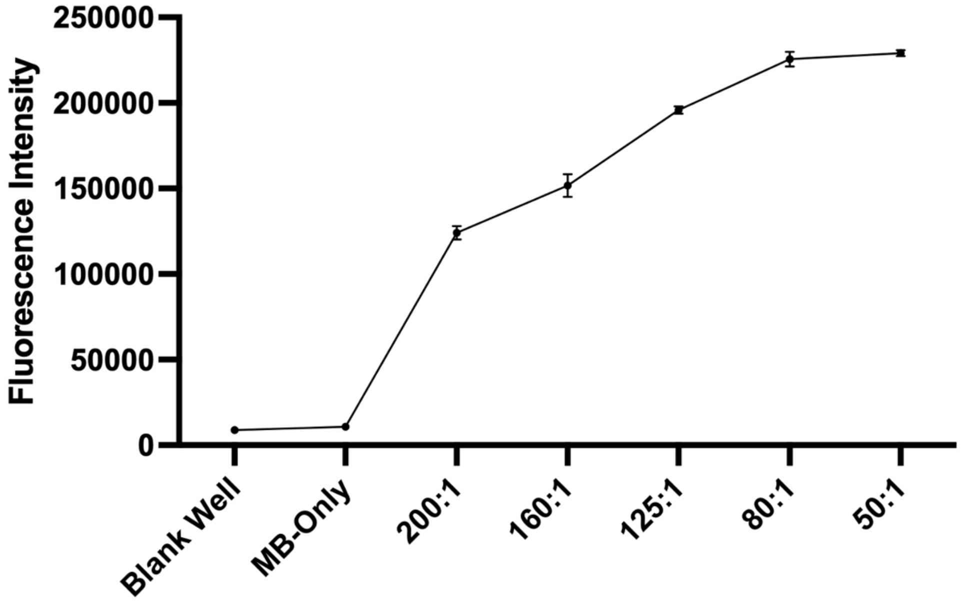


**Supplementary Figure 6.** FI plot of MB-strep-Ab conjugates prepared with different MB:Ab mass ratios, including controls without Ab and blank well readings.

### Supplementary Discussion 7: Magnetic Field Simulation

For designing the MagChip, to understand the influence of the magnetic field distribution on the MB pattern formation within the POC wells, we conducted simulations of the magnetic field distribution using COMSOL Multiphysics. **Supplementary Figure 7** illustrates the simulated magnetic field profiles under two different configurations: one is a central pair of magnets surrounded by additional magnets, and the other is the same central pair of magnets without any surrounding magnets. In our SIMAIS biosensor design, although only two magnets are positioned directly beneath the POC wells and are primarily responsible for inducing the MB pattern formation, the inclusion of surrounding magnets plays a crucial role in achieving a uniform magnetic field across the wells.

The primary function of the central magnets is to generate a magnetic field that influences the MBs within the wells, causing them to aggregate and form patterns indicative of target protein concentrations. However, when only the central magnets are present, the magnetic field lines exhibit significant distortion at the edges of the magnets. This distorted magnetic field leads to inconsistencies in MB behaviour and pattern formation. By incorporating surrounding magnets with the same polarity orientation as the central magnets, the magnetic field lines become non-distorted and similar within the region of interest - wells in the POC chip. The surrounding magnets effectively reduce edge effects and magnetic field gradients by providing a more homogeneous magnetic environment. This uniformity results in a more reliable and reproducible pattern formation.

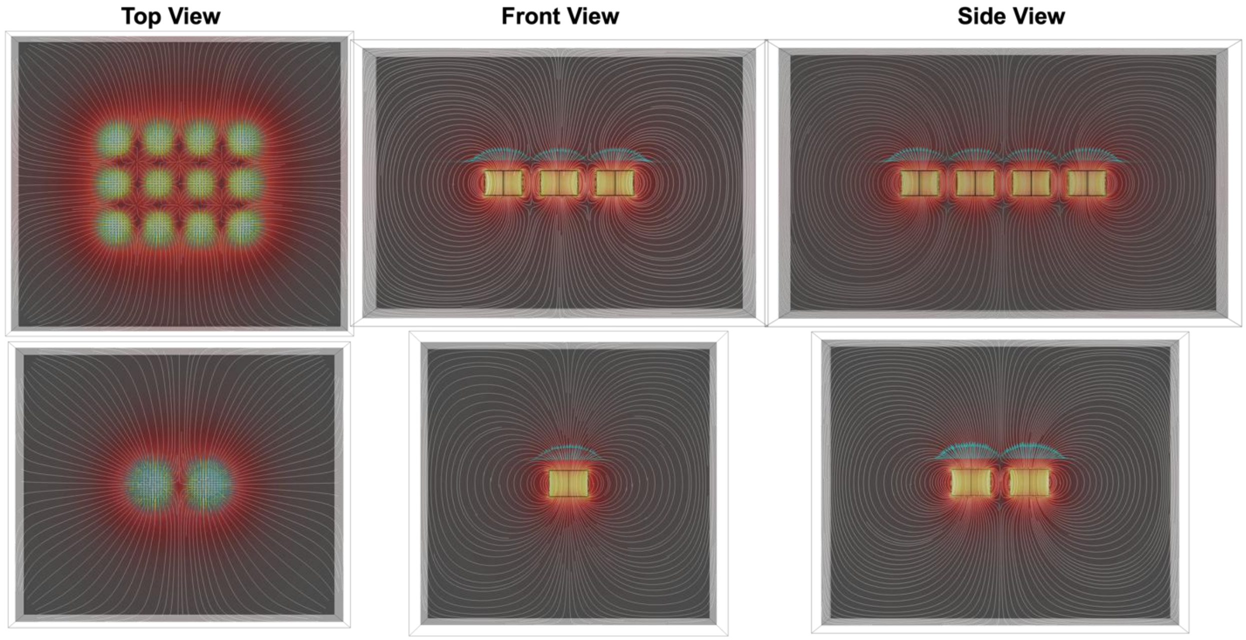


**Supplementary Figure 7.** Magnetic field simulations illustrating the effect of surrounding magnets on field distribution within the POC chip wells (Top panel: a central pair of magnets surrounded by additional magnets of the same type and orientation; Bottom panel: only the central pair of magnets without any surrounding magnets). All magnets used are 6mm (diameter) x 4mm (height) and graded as N52, with the north pole facing upwards from the front view.

### Supplementary Videos

**Supplementary Video 1.** Conceptual introduction to the SIMAIS biosensing platform, showing its inspiration basis in swarm intelligence and highlighting its key features and advantages.

**Supplementary Video 2.** Top-view experiment video recording of MBs pattern formation under a magnetic field, comparing bare MBs, antibody-conjugated MBs without target antigen, and antibody-conjugated MBs with target antigen.

**Supplementary Video 3.** Schematic animation illustrating the mechanism of MBs aggregation and pattern formation.

**Supplementary Video 4.** Demonstration of the SIMAIS biosensing platform in a point-of-care setting, showing integration with a smartphone app.
